# Effects of Unleaded Petroleum on the Macrophage Aggregates (MA) formation in Red King tilapia (*Oreochromis sp.*) Fingerlings

**DOI:** 10.1101/044537

**Authors:** Ancel Jeff G. Beso, Veronica Y. Candelaria, Jennifer G. Dela Cruz, Margie S. Tolentino, Anna Danica C Tameta, Allen A. Espinosa

## Abstract

The Philippines is one of the major producers of tilapia, the most cultured fish and widely consumed in the world. Although fishes in general is said to be adapted to various stressful conditions, the effect on several cellular immune parameters may be of interest to determine the capacity of the organism to withstand stressors. In this paper, the effect of unleaded petroleum on the splenic macrophage aggregate (MA) formation was studied. This was done to have an overview of the immune response of Tilapia or fishes in general when an oil spill, which almost occur annually at different parts of the world, happen. Histological analysis assessed the area occupied by splenic MA 24 hours after introduction of unleaded petroleum to the aquatic system. To determine whether Mabuhay balls, a technology that claims to be beneficial in terms of improving water quality, was added to one tank (T1) to be able to compare it with another tank (T2). There is a strong statistically significant difference between the groups at day1 (p=0.000) opposite the result of day 6 (p=0.155). Thus, unleaded petroleum increased MA formation, a sign that may indicate a high immune activity as an initial positive response to stress. Mabuhay ball have lessen the mortality but has no effect on splenic MA formation.

## INTRODUCTION

Tilapia is the second most cultured fish after carps (Nia, Xin Zhou, et al. 2013). Tilapia is one of the most important commercial fresh water cultivated worldwide thanks to its fast growth rate in warm waters, it’s tolerance to adverse conditions, relatively low production cost, meat quality and flavor, high protein content and consumer reference (Tellez-Banuelos et al. 2008)

In the Philippines, oil spill last happened in August 9, 2013. Five hundred thousand litters of petroleum was accidentally scattered for almost 300sqm. along Manila Bay (newsinfo.inquirer.net, 2013). Due to a competent immune system, fish will often try to avoid pollutants when they can, but maybe trapped or compelled by reproductive instincts to swim through oil towards their mating grounds. After a sudden, large release, those fish that can swim away might be exposed to significant quantity of oil before escaping (Davis et al. 2002). Fish and mammals; immune system are more similar than different in terms of their physiological and biological characteristics.( WOLF et al. 2005)

Macrophage Aggregates (MAs) occurs in the spleen, head kidney and liver of most teleosts, it is splenic MA’s that best serve as a reliable histopathological bio indicator of fish health and environment degredation (Fournie et al. 2001).

The spleen is usually a solitary, dark red in the peritoneal cavity adjacent to the wall gut. The same basic element as in higher vertebrate are typically present: blood vessels, red and white pulps and ellipsoids. As fish have no lymph nodes, the spleen alone plays an essential role in antigen trapping. The spleen is the major peripheral lymphoid organ (Mahabady, et al. 2012). The objectives of this research is to assess the effect of oil spill to Red King Tilapia fingerlings and to assess the capability of Red King Tilapia to adopt to a polluted environment. In addition, the effect of Mabuhay balls on mortality and MA formation was also assessed based on the hypothesis that Mabuhay ball can improve water quality which equates to improving fish health in general.

## METHODOLOGY

Approximately 50 mixed-sex of approximately two weeks clinically healthy Red King Tilapiawere obtained from a distributor in Novaliches, Quezon City. The fish were said to be reared from a fish farm in Laguna. Upon purchase all fish were immediately transferred to the CLAB, Colegio de San Juan de Letran.

### Sampling 1

Acclimatization was done for 10 days and during this period; fish were fed once a day to satiation. After acclimatization the fish was randomly distributed in three aquaria labelled C, T1, T2. The average standard length and weight per treatment group were as follows: T1(46.33±8; 3.06±3) T2(52.84±10; 3.89±0.1) C(48.5±1; 2.49±1). After 10 days of acclimatization, petroleum (unleaded, Petron) was added to obtain a concentration 0.5ml per litre of water for T1 and T2, No petroleum was added to C (negative Control). After 24 hours of exposure to petroleum, 3 fish was collected from T1 and T2 and 2 fish from C. Ice was used as an anaesthetic to immobilize the fish.

Dissection of fish was done by cutting from the anus to the posterior margin of the operculum. Spleen from each fish were excised and placed in 10% formalin. These samples were processed for routine histological procedure at PKDF (Philippine Kidney Dialysis Foundation) located at Roces Avenue, Quezon City. All spleen samples were stained with H&E.

### Sampling 2

After 6 days of exposing the fish to petroleum wherein T1 is with bukashi ball and T2 is stays as the same. Ice was used as an anaesthetic to immobilize the fish. Dissection of fish was done by cutting from the anus to the posterior margin of the operculum. Spleen from each fish were excised and placed in 10% formalin. These samples were processed for routine histological procedure at PKDF (Philippine Kidney Dialysis Foundation) located at Roces Avenue, Quezon City. All spleen samples were stained with H&E.

Using a microscope, area occupied by MA was observed at LPO using a light microscope. Iphone5 HDR camera was used to capture the images and five MAs per slide were photographed in random fields of view and were measured using ImageJ. One way ANOVA was used to determine statistical difference between treatment groups.

## RESULTS

Tilapia are wildly cultured because they breed easily and could adapt to different environmental conditions. This is one of the most significant species for fish farming and for biological studies as well. It is also represents a good study model for immune toxicity studies (Tellez-Banuelo et al. 2008).

In sampling 1 the splenic MA is statistically significant between treatment groups at p= 0.000. At day 6, high mortality was observed in T2 (without mabuhay ball) that prompt the researchers to terminate the experiment immediately. However, statistical analysis showed that there is no significant difference in splenic MA formation (p=0.155.

## DISCUSSION

Compared to a parallel experiment, MA formation in spleen as a response to stress was very pronounced compared to head kidney. This indicated that spleen, in this case, was more responsive to sudden stress experienced in the aquatic system. However, it may also be accounted that the spleenic MA might be already present even before the said experiment and it just enhances the MAs (Montero et al, 1998 in Tameta, 2010 graduate thesis).

Splenic MA were observed and was measured using Image J and the splenic MA was compared between groups. Movement of contaminants into the flesh of fish requires that they are in water so that they may rapidly pass via the gills into the bloodstream, thence to circulate throughout the tissues. The concentration of the oil components in muscle of fish exposed to water-solubilised fractions of fuel oil increased rapidly, reaching a maximum within approximately 1 hour ( Davis,et al. 2002). The fish was exposed to petroleum 24 hours and dissected 3 fishes from T1 and T2 and 2 fish from C, and had H&E stain on their spleen to use the macrophages, Where Day 1 (Figure 1) is 0.000 p value and with week 1 is 0.155 p value and the outcome of the results that we can see is that the splenic MAs still occurred in week 1 however the mortality was lessen due to Mabuhay balls yet doesn’t give any further significant reduction on splenic MAs.

**Fig. 1.**
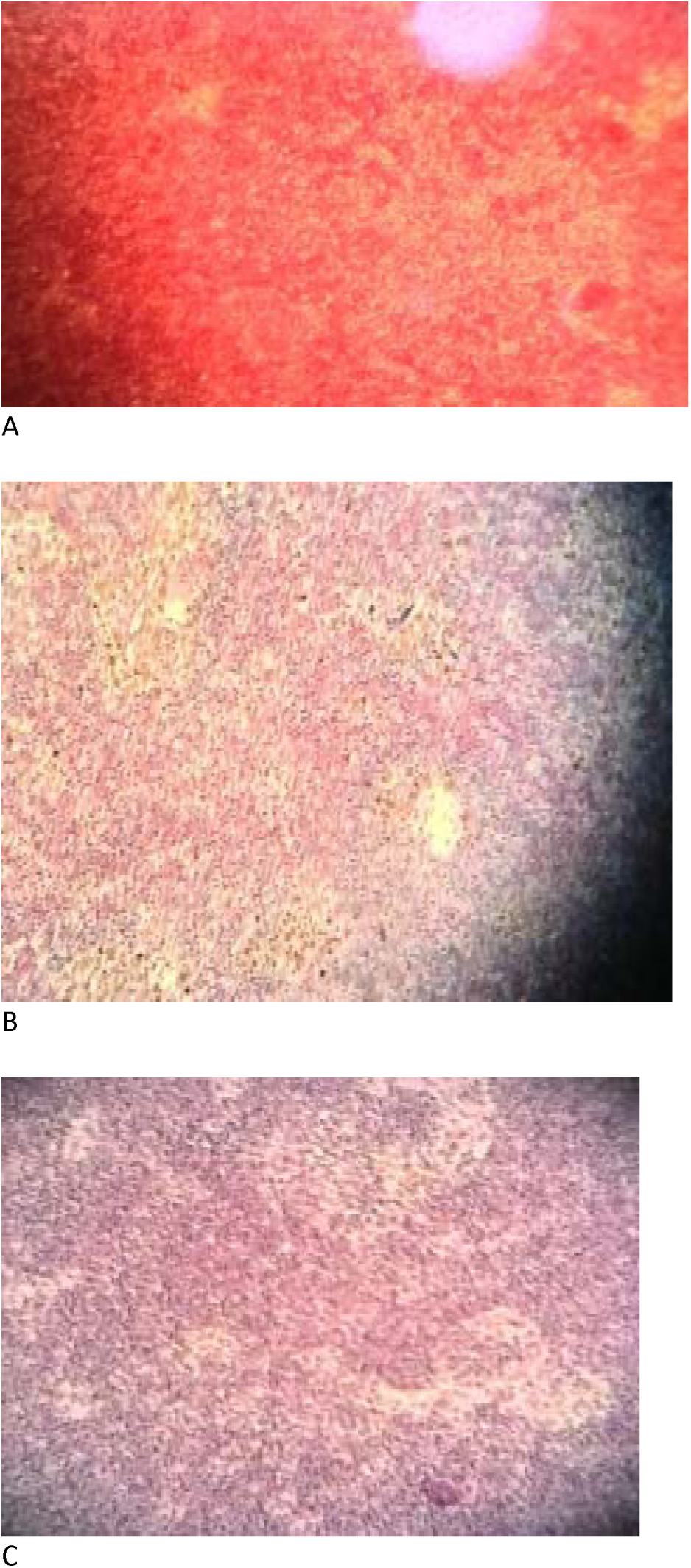
Splenic MA at day 1: A (T1), B (T2), and C (control) showing significant difference in area occupied by MA. Brown color indicates MA formation.

The spleen is the major organ in terms of immunological responses however the head kidney is the first line of defense in fishes (Tameta, 2013. Graduate thesis). The formation of MA in the head kidney of sampling 1 p= 0.500 and in week 1 is p= 0.000 (Ligutan et al. 2013. Unpublished research) compare with spleen with sampling 1 p= 0.000 and in sampling 2 p= 0.155 this might be because the spleen is the major organ to response in the immune system and the formation of MA might also be prior to the experiment. However at day 6 (figure 2), the fish might be able to adapt to the stressful condition thus, not showing any significant difference in MA formation. This result is similar to the result of Harford et al. 2005 (in Tellez-Banuelos et al. 2008) that the formation of MA of spleen and head kidney increases after exposing the juvenile tilapia in endosulfan(pesticide) for 21 days wherein the adaptation to the polluted environment is statistically significant.

**Fig. 2.**
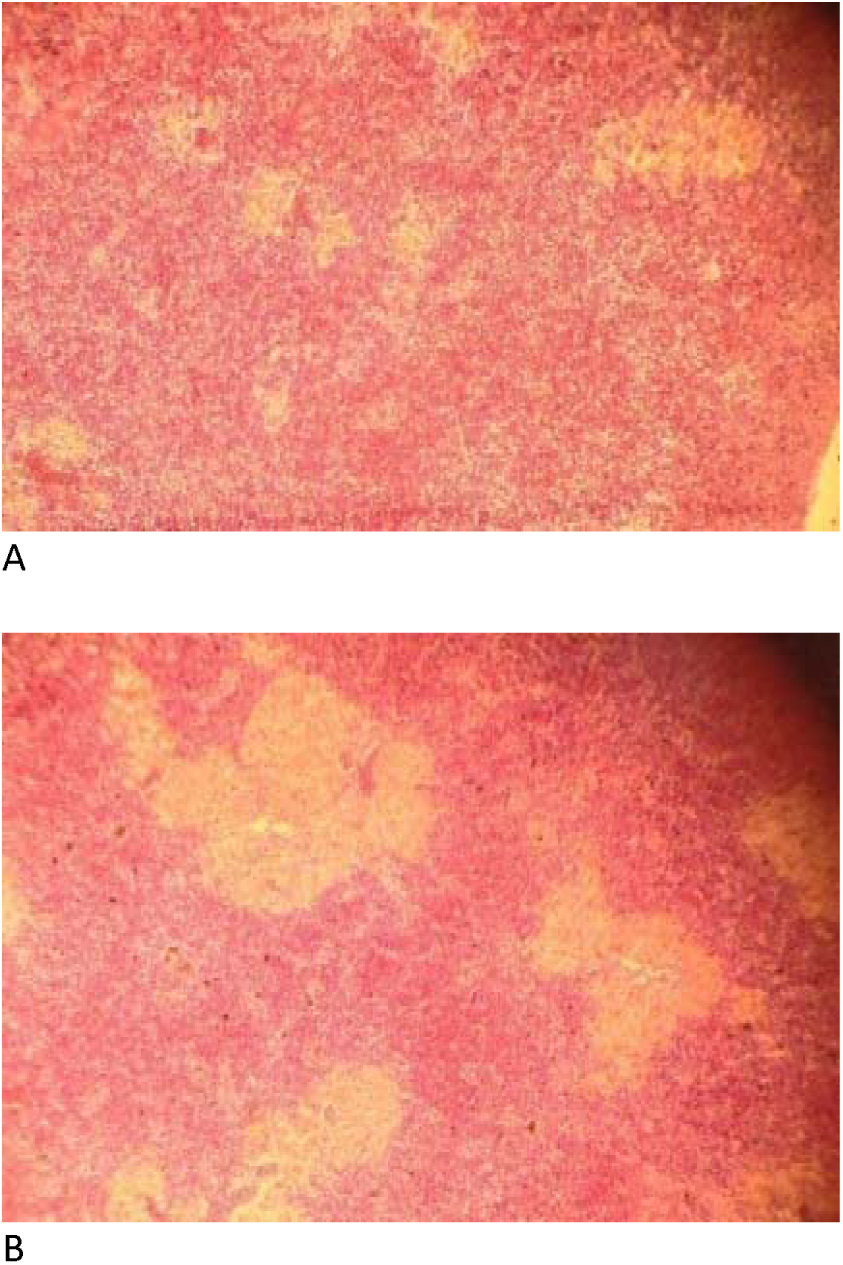
Splenic MA at day 6: A(T1) with Mabuhay balls and B (T2). Brown color indicates MA formation.

## Conclusion

Based on the experiment and it’s result the petroleum gives a very significant formation of spleenic MAs, in this way we can assume that the sampling 1 of the experiment showed us the negative effect of exposing the fish to petroleum after more than 24 hrs. Addition of Mabuhay ball in T1 might be assumed to be beneficial since high mortality was observed in T2 (without Mabuhay ball), however, it did not affect MA formation. The significance of the mabuhay ball can’t be stated because it possibly needed further information and study. It is also recommended that more samples will be used in future research to strengthen statistical validity. Also, special staining such as Perl’s Prussian blue may be used to have a better image and comparison on MA formation.

